# Revised 16S rRNA V4 hypervariable region targeting primers enhance detection of *Patescibacteria* and other lineages across diverse environments

**DOI:** 10.1101/2025.11.26.690684

**Authors:** Huifeng Hu, Clemens Karwautz, Kalina Duszka, Thomas Karner, Isabella C. Wagner, Christoph Grander, Wilhelm Grander, Laura Steinwidder, Lucilla Boito, Viktor Van de Velde, Marijn Bauters, Pascal Boeckx, David Seki, Bettina Glasl, Stefan Thiele, Hannes Schmidt, Joana Séneca, Michael Wagner, Petra Pjevac

**Affiliations:** Centre for Microbiology and Environmental Systems Science, University of Vienna. Djerassiplatz 1, 1030 Vienna, Austria; Doctoral School in Microbiology and Environmental Science, University of Vienna. Universitätsring 1, 1010 Vienna, Austria; Department of Functional and Evolutionary Ecology, Faculty of Life Sciences, University of Vienna, Djerassiplatz 1, 1030 Vienna, Austria; Department of Nutritional Sciences, Faculty of Lifesciences, University of Vienna, Josef-Holaubek-Platz 2, 1090 Vienna, Austria; Centre for Animal Nutrition and Welfare, University of Veterinary Medicine, Veterinärplatz 1, 1210 Vienna, Austria; Faculty of Psychology, University of Vienna, Liebiggasse 5, 1010 Vienna, Austria; Department of Internal Medicine I, Gastroenterology, Hepatology, Endocrinology, and Metabolism, Medical University of Innsbruck, Innrain 52, 6020 Innsbruck, Austria; Department of Internal Medicine, Hall State Hospital, Milserstrasse 10, 6060 Hall in Tirol, Austria; Biobased Sustainability Engineering (SUSTAIN), Department of Bioscience Engineering, University of Antwerp, Antwerp 2020, Belgium; Department of Environment, Faculty of Bioscience Engineering, Ghent University, Coupure Links 653, 9000 Gent, Belgium; Department of Green Chemistry and Technology, Faculty of Bioscience Engineering, Ghent University, Coupure Links 653, 9000 Gent, Belgium; Department of Laboratory Medicine, Medical University of Vienna. Währinger Gürtel 18-20, 1090 Vienna, Austria; Joint Microbiome Facility of the Medical University of Vienna and the University of Vienna. Djerassiplatz 1, 1030 Vienna, Austria; Center for Microbial Communities, Department of Chemistry and Bioscience, Aalborg University. Fredrik Bajers Vej 7H, 9220 Aalborg, Denmark; Environment and Climate Hub (ECH), University of Vienna, Augasse 2-6, 1090 Vienna, Austria

**Keywords:** 16S rRNA gene amplicon sequencing, primer modification, Patescibacteria

## Abstract

Primer bias in 16S rRNA gene amplicon sequencing can distort microbial diversity estimates by underrepresenting key taxa. We introduce a modified primer pair (V4-EXT) targeting the hypervariable V4 region of bacterial and archaeal 16S rRNA genes, with improved *in silico* taxonomic inclusivity. To benchmark performance, we analyzed 938 samples from terrestrial, aquatic, and host-associated habitats, comparing microbial community profiles derived with V4-EXT and the currently most widely used V4-targeted primers. V4-EXT substantially improved the detection of *Patescibacteria* and other underrepresented lineages, such as *Chloroflexota* and *Iainarchaeota*, while enhancing recovery of novel amplicon sequence variants across sample types. Overall, V4-EXT provides broader taxonomic coverage and more inclusive microbial community profiles, particularly in high-diversity ecosystems such as groundwater and soils. We propose V4-EXT as a robust successor for comprehensive microbial community analysis across diverse habitats.

## Background

Gene-targeted amplicon sequencing remains the most widely used sequencing-based technique in microbial ecology and microbiome research. It enables low-cost, high-throughput analyses of microbial community composition and potential metabolic functions in diverse environments by targeting, amplifying, and sequencing phylogenetic or functional marker genes [1–6]. Despite its utility, gene-targeted amplicon sequencing has several limitations. A major concern is primer bias, as mismatches between primers and target sequences can result in selective amplification and potentially skew diversity estimates, community composition profiles, and ecological interpretations [7,8].

Most studies analyzing microbial communities use broad-range primers that target the 16S rRNA phylogenetic marker gene. Since the widespread implementation of short-read sequencing, primers targeting conserved regions around short, hypervariable stretches (mainly V1-V2, V3-V4, V4, and V9), generating 200-350 bp amplicons, have become commonplace. The most commonly used amongst these are the bacterial and archaeal 16S rRNA V4 targeted primers 515F [7] and 806R [8], used by the Earth Microbiome Project and hereafter referred to as “V4-EMP” [1]. In a recent study, we demonstrated that a modification of these V4 primers significantly improved the detection of *Patescibacteria*. By applying the optimized primer (hereafter referred to as “V4-CPR”) to a global collection of wastewater treatment plant (WWTP) samples, we revealed that *Patescibacteria* constitute a substantial and previously largely overlooked fraction of WWTP microbial communities [9]. However, V4-CPR has only been experimentally evaluated with WWTP samples and has shown limited *in silico* coverage of some archaeal lineages that are typically low in abundance in WWTPs. In the present study, we further increased the taxonomic inclusivity of the primers by improving coverage of archaea via reintroducing a single-base degeneracy in the forward primer (at position 523, A to M), while maintaining improved *Patescibacteria* detection.

We evaluated the performance of the revised 16S rRNA V4 primer set, hereafter referred to as “V4 extended” or “V4-EXT”, by applying it to 938 samples from various sources, including soil, freshwater, and marine environments, as well as human and animal stool. We analyzed differences in community alpha diversity, community taxonomic profiles, and detected sequence novelty between datasets generated with the V4-EXT and V4-EMP primers. Our results demonstrate that the V4-EXT primers offer an expanded taxonomic detection range and improved sensitivity for *Patescibacteria* and other taxa. This results in more inclusive microbial diversity profiles, especially when applied to soil, freshwater, and ocean samples.

Based on these findings we recommend using the V4-EXT instead of V4-EMP in future studies, as it enables more comprehensive microbial community profiling across different habitats.

## Results

### Primer modifications enable expanded in silico coverage of the archaeal and bacterial 16S rRNA gene V4 region

We evaluated the *in silico* coverage of three V4 primer sets against bacterial and archaeal sequences, using the most recent release of the SILVA database (r138.2 SSURef NR99) [10] as a reference. We compared without mismatch and one mismatch coverage for the V4 primer pair 515F [7] and 806R [8] used by the Earth Microbiome Project [1] (V4-EMP), the V4 modified primer pair optimized for *Patescibacteria*, previously referred to as Candidate Phyla Radiation [9] (V4-CPR); and the V4 primer pair further modified for extended taxonomic coverage, which was developed in this study (V4-EXT).

Our analysis showed that V4-EMP primers provided robust *in silico* coverage of most bacterial and archaeal lineages but only captured 19.6% (69.6% with one mismatch) of *Patescibacteria* and only 56.7% (89.5% with one mismatch) of *Chloroflexota* sequences (Table S1). The V4-CPR primers achieved an 88.9% (98.5% with one mismatch) and 76.0% (95.2% with one mismatch) *in silico* coverage of *Patescibacteria* and *Chloroflexota* 16S rRNA genes, respectively. However, they exhibited limited without mismatch coverage across most archaeal phyla (Table S1). The V4-EXT primers offered a more balanced performance, improving coverage across all bacterial and archaeal phyla, compared to the V4-EMP and V4-CPR primers (Table S1). Notably, the V4-EXT primers significantly improved the *in silico* coverage of archaeal groups poorly covered by the V4-CPR primer, such as the *Iainarchaeota* and *Altiarchaeota* phyla (Table S1).

### In situ evaluation of V4-EXT primers across various environmental samples

We experimentally evaluated and compared the performance of the V4-EMP and V4-EXT primers. We generated and analyzed amplicon datasets from a total of 938 samples collected from four distinct habitats, including 361 soil samples, 155 samples from ground and freshwater, nine samples from sponges, 160 samples from surface ocean water, and 253 fecal samples from humans, mice, and cows. These samples originated from 12 independent studies, with sample collection and DNA extraction methods selected according to sample type (see Table S2; see Methods for further detail).

Co-amplification of eukaryotic 18S rRNA gene V4 regions was only detectable in soil and freshwater samples at negligible frequencies with both primer pairs (Table S3). Co-amplification of chloroplast 16S rRNA gene sequences depended on the environment and differed significantly between datasets. Yet, the proportions of co-amplified chloroplast 16S rRNA genes were similar between primer pairs and were most pronounced in ocean samples, where 15.71% and 17.66% of reads were classified as chloroplast in datasets generated with the V4-EMP and V4-EXT primers, respectively (Table S3). In freshwater samples, 6.29% and 5.14% of reads were identified as chloroplast-derived in the V4-EMP and V4-EXT datasets, respectively. Chloroplast-derived reads were also detected in soil samples but at much lower levels (1%-1.5%), and were nearly absent in gut samples (0.02%) with both primer pairs (Table S3). In contrast, mitochondrial 16S rRNA gene sequences were consistently co-amplified at a significantly higher proportion with the V4-EXT primers (Table S3). In soil samples, the V4-EXT primers resulted in 8.34% of reads being classified as mitochondrial 16S rRNA genes, nearly three times the proportion obtained with the V4-EMP primers (2.83%). A similar pattern was observed in ocean surface water and freshwater samples, where the V4-EXT primers yielded three to five times more mitochondrial reads. In fecal samples, mitochondrial sequences were detected at low levels with both primer pairs (<0.1%) (Table S3). For all downstream analyses, reads classified as chloroplast, mitochondrial, eukaryotic, or unclassified at the phylum level were removed from the datasets.

For comparisons of alpha diversity (richness and evenness), each sample in both the V4-EMP and V4-EXT datasets was rarefied to 5,000 observations. Significantly higher observed amplicon sequence variant (ASV) counts and Shannon diversity were observed in the V4-EMP dataset from soil samples (Figure 1A, 1B). In contrast, the V4-EXT primers resulted in significantly higher observed ASV counts and elevated Shannon diversity (Figure 1A, 1B) in freshwater and groundwater datasets, while only Shannon diversity was found to be significantly higher when using the V4-EMP primers in the open ocean surface water dataset (Figure 1B). The V4-EXT primers detected a significantly higher number of observed ASVs in human stool samples, whereas they detected significantly fewer ASVs than the V4-EMP in cow stool samples (Figure 1A). Alpha diversity metrics did not differ significantly when the two primer pairs were applied to mouse stool samples (Figure 1A, 1B). Overall, regression analyses across all samples showed comparable alpha diversity was recovered by the two primer pairs based on both observed ASV counts and the Shannon diversity index (Figure 1C, 1D).

**Figure 1.**
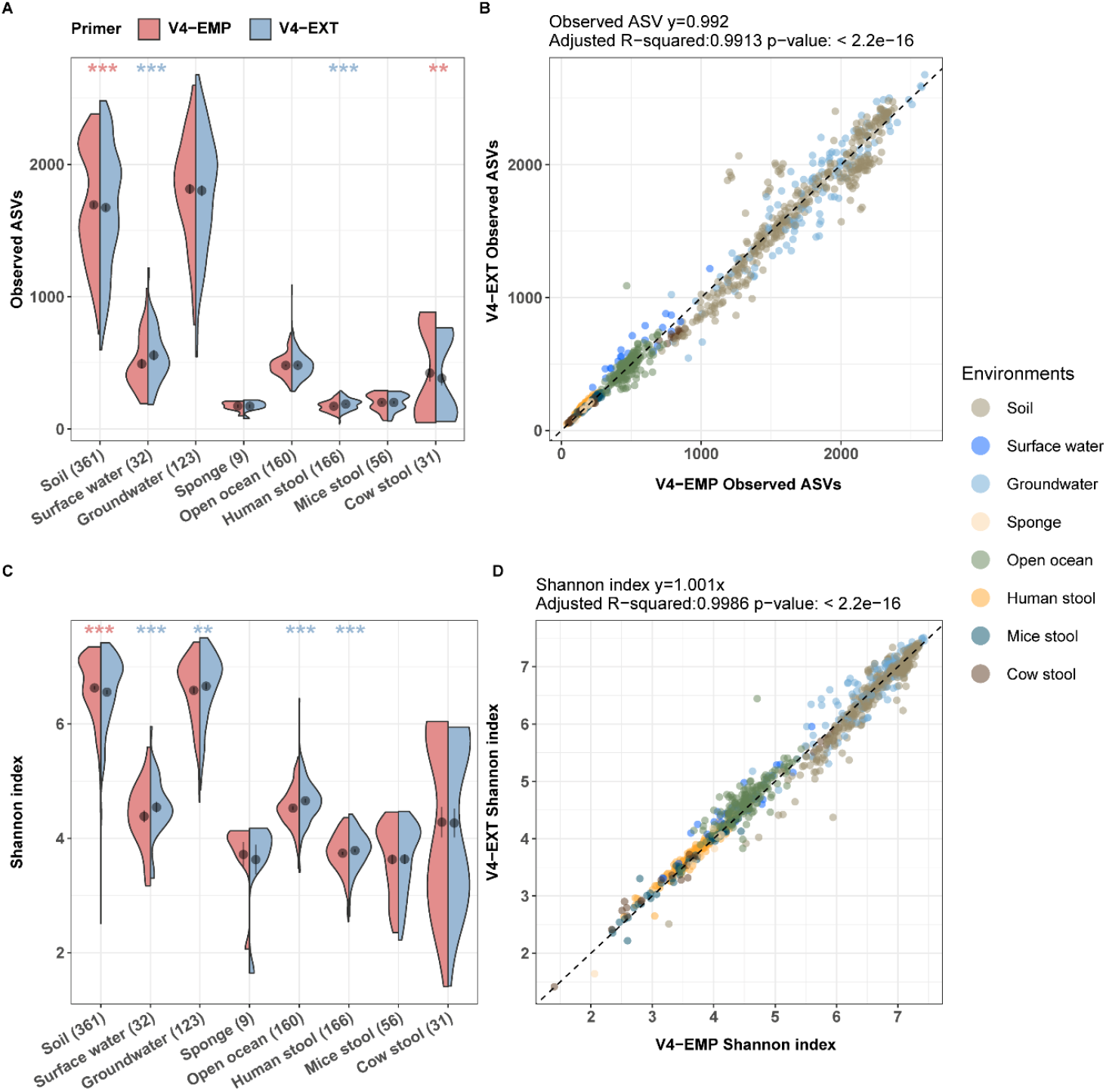
Comparison of (A) observed ASV counts and (B) Shannon index in different datasets generated using the V4-EMP (red) and V4-EXT (blue) primers. The number of samples (in parentheses) and the source environment for each dataset are shown on the x-axis. Points and error bars, respectively, represent the mean and standard error (SE) for each dataset. Statistical significance of differences between the datasets was tested by paired Wilcoxon signed-rank tests. *: p < 0.05; **: p < 0.01; ***: p < 0.001. The significance indicators are colored in red if the V4-EMP primer dataset has a higher median value and in blue if the V4-EXT primer dataset has a higher median value. Regression analysis of (C) observed ASV counts and (D) Shannon index values between datasets generated using the V4-EMP and the V4-EXT primer pairs. The correlation coefficient, R_2_ value and p-value are indicated in the title of each panel. The theoretical 1:1 line is indicated as a dashed line.

### Relative abundance difference at phylum level across different environments

Next, we compared phylum-level differential abundance patterns between datasets generated by the two primer pairs. When using the V4-EXT primers, the relative abundance of *Patescibacteria* was more than twice as high in all sample types except human stool samples, where their average abundance was below 0.1% (Figure 2A,2B, Table S4). For example, *Patescibacteria* accounted for on average 39% of all reads in groundwater samples from the V4-EXT dataset, but only for 6.4% in the V4-EMP dataset. In surface freshwater samples, *Patescibacteria* were less dominant, but the V4-EXT primers detected 4.4% reads while the V4-EMP primer detected only 0.3%. In soil samples, the V4-EXT dataset contained on average 13.1% *Paterscibacteria* reads, while the V4-EMP dataset contained only 0.8% (Figure 2B, Table S4). Interestingly, in ocean surface water samples, where *Patescibacteria* were rare regardless of primer choice, *Bacterodiota* were over 10% more abundant in the V4-EXT dataset (39.2%) than the V4-EMP dataset (28.6%) (Figure 2B, Table S4). Similarly, *Planctomycetota* accounted for 7% in V4-EXT datasets from ocean surface samples, but only for 2.6% in the matching V4-EMP dataset. Furthermore, *Chloroflexota* were generally more abundant in the V4-EXT datasets, especially in ocean, soil, and sponge samples. The V4-EXT primers additionally resulted in significantly higher relative abundances detected for several other phyla in specific environments (Figure 2A, Table S4): *Poribacteria* and *Iainarchaeota* in groundwater samples; *Cyanobacteria*, Latescibacterota, and GAL15 in soil samples; *Planctomycetota* and *Chloroflexi* in ocean surface water samples; and *Planctomycetota, Elusimicrobiota*, and *Chloroflexi* in cow stool samples. Conversely, the V4-EMP datasets contained statistically significantly higher relative abundances of five archaeal phyla (*Thermoplasmatota, Nanoarchaeota, Halobacterota, Euryarchaeota*, and *Aenigmarchaeota*) across all sample types (Figure 2A, Table S4). Among these, *Thermoplasmatota* were particularly abundant (19%) in the open ocean surface water sample, and *Nanoarchaeota* (35.1%) in the groundwater sample datasets generated with V4-EMP primers, but accounted for only 6.5% and 10.5%, respectively, in V4-EXT datasets (Figure 2A, Table S4). The remaining archaeal phyla were comparatively rare in both datasets from the samples analyzed in this study. Furthermore, seven bacterial phyla (*Elusimicrobiota, Nitrospirota, Myxococcota, Firmicutes*, FCPU426, *Fibrobacterota*, and RCP2−54) were relatively more abundant in the V4-EMP soil datasets, while one phylum was more abundant in the V4-EMP dataset from groundwater (*Nitrospirota*), human stool (*Verrucomicrobiota*), and sponge (*Bacteroidota*) samples, respectively. Although not being statistically significantly different, *Firmicutes*-related sequences were notably more abundant in the V4-EMP (16.9%) than the V4-EXT (9.3%) soil dataset, while *Cyanobacteria* and *Nitrospirota* related reads were double as abundant in the V4-EMP dataset (11.5% and 11.2%) than the V4-EXT dataset (6.2% and 5.0%) from sponge samples.

**Figure 2.**
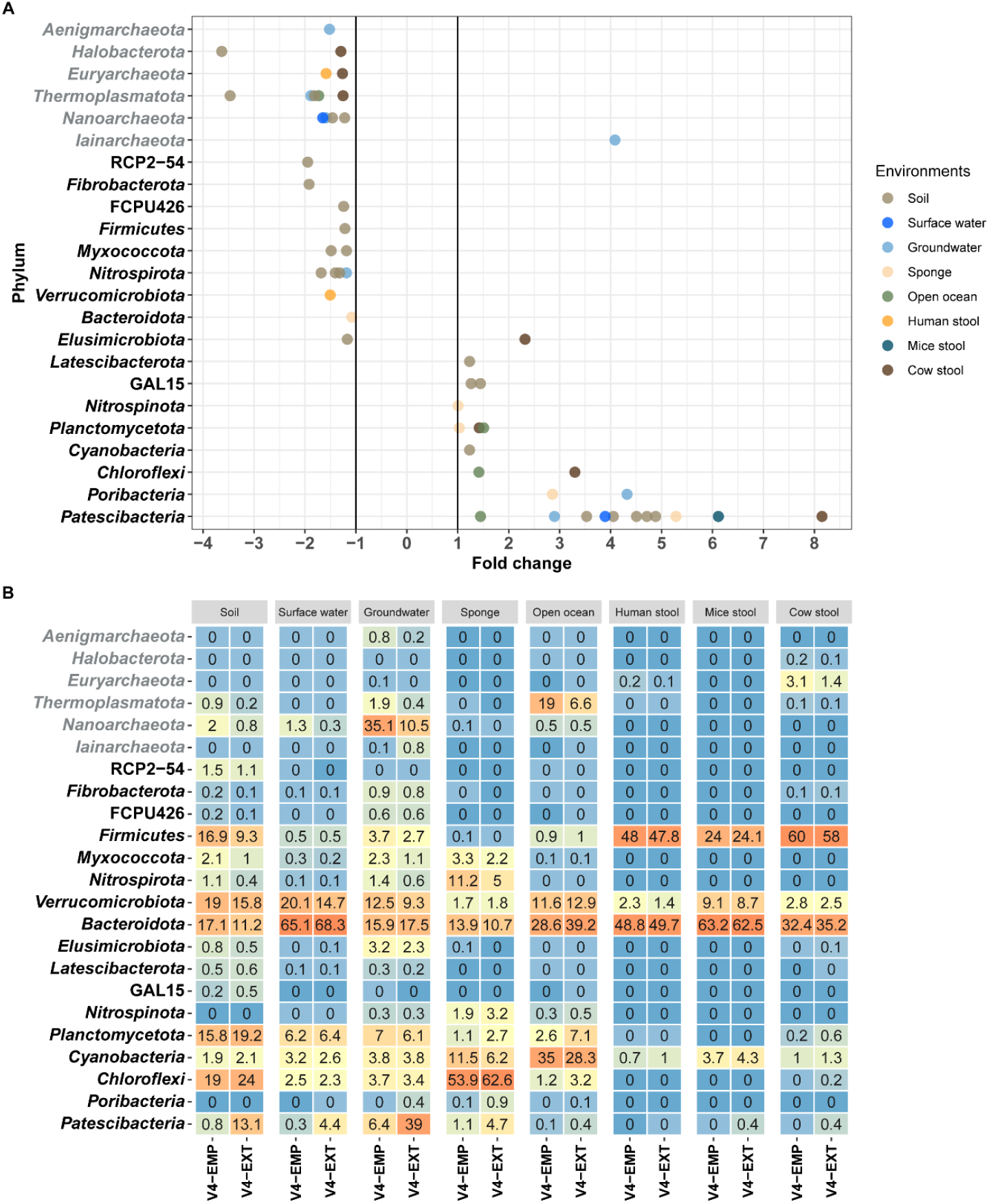
(A) Fold changes in the relative abundances of each phylum in datasets colored by source environment. Each dot represents a research project and dots are colored by different source habitats. Phyla with a mean relative abundance of >0.1% for which mean relative abundance differences between datasets were above twofold are depicted. Fold change >1 indicates the phylum has a higher mean relative abundance in the V4-EXT dataset and fold change <-1 indicates the phylum has a higher mean relative abundance in the V4-EMP dataset. (B) Phyla level average relative abundance heatmap across different types of samples grouped by source environment, generated using the V4-EXT versus the V4-EMP primers. Phyla with a mean relative abundance of >0.1% in either dataset are presented. Archaeal phyla are shown in grey font, and bacteria phyla are shown in black font.

As the relative abundance of *Patescibacteria* was significantly higher in all V4-EXT sample types except human stool samples, pronounced changes in their relative abundances could have masked or inflated shifts in the relative abundance of other phyla. To better evaluate the effect of primer choice on the residual community, we compared the relative abundances of phyla between datasets after removing reads classified as *Patescibacteria*. As expected, the differences in the relative abundance of other phyla between the V4-EXT and V4-EMP datasets from soil and groundwater samples became less pronounced, as *Patescibacteria* were highly abundant in the V4-EXT datasets from these samples (Figure S1A,B, Table S4). In groundwater samples, the relative abundance of archaea increased, with 22.4% and 38.4% of reads classified as *Nanoarchaeota* in the V4-EXT and V4-EMP datasets, respectively. Yet, *Nanoarchaeota* and other rare archaeal phyla still had a higher relative abundance in V4-EMP datasets. However, the relative abundance of *Iainarchaeota* was approximately four times higher in the V4-EXT dataset (Figure S1A). While these results might suggest a potential primer bias affecting specific archaeal phyla, the overall low relative abundance of these phyla, even in the V4-EMP groundwater dataset, must be noted. Such low relative abundance may exaggerate differences detected in differential abundance analyses (Figure S2), as the differentials are more strongly affected by the improved detection and thus increased relative abundance of other taxa. Notably, the relative abundance of numerous bacterial phyla, including *Bacteroidota, Planctomycetota, Chloroflexi*, and *Cyanobacteria*, was correspondingly substantially higher in the V4-EXT dataset. A similar trend was observed in soil samples after excluding *Patescibacteria* reads, with a higher relative abundance of *Planctomycetota* and *Chloroflexi* in the V4-EXT dataset, while *Verrucomicrobiota, Bacteroidota*, and *Firmicutes* remained relatively more abundant in the V4-EMP dataset (Figure S1).

### More novel ASVs are observed in V4-EXT datasets

Interpreting 16S rRNA gene amplicon data relies heavily on databases that provide the necessary taxonomic, functional, and ecological context for ASVs or operational taxonomic units (OTUs) generated from raw sequences. ASVs and OTUs without close relatives in these databases likely represent novel biological entities. Thus, we focused specifically on such novel sequence variants generated by the V4-EMP and V4-EXT primer pairs.

First, we compared the fraction of novel ASVs, lacking high-identity matches (<99% identity) [11] or genus-level matches (<94.5% identity) [12] in the SILVA database. The V4-EXT primers detected significantly more novel ASVs at both thresholds across all sample types except cow stool samples, for which both primer pairs yielded similar proportions of novel ASVs (Figure 3A). Groundwater samples exhibited the highest proportion of novel ASVs with both primer pairs. Nearly half of the ASVs obtained from groundwater samples by the V4-EXT primers (56.5% ± 12.0%, mean ± SD) lacked high-identity matches in the SILVA database, and 23.4% ± 9.3% of these ASVs could not be assigned to a genus. Using the V4-EMP primers on the same samples yielded 51.5 ± 10.7% ASVs without high-identity matches in the SILVA database (Figure 3A,3B). As expected, stool samples from humans, mice, and cows showed the lowest levels of novelty (on average 4.6% ± 4.1% and 8.4% ± 6.0% in the V4-EMP and V4-EXT datasets, respectively), confirming that these microbial communities are comparatively well characterized. Across all samples, cumulatively, 33.0% of ASVs from the V4-EXT dataset and 30.7% from the V4-EMP dataset lacked high-identity matches. At the genus level, 11.6% (V4-EXT) and 8.6% (V4-EMP) of ASVs could not be assigned across all samples cumulatively.

**Figure 3.**
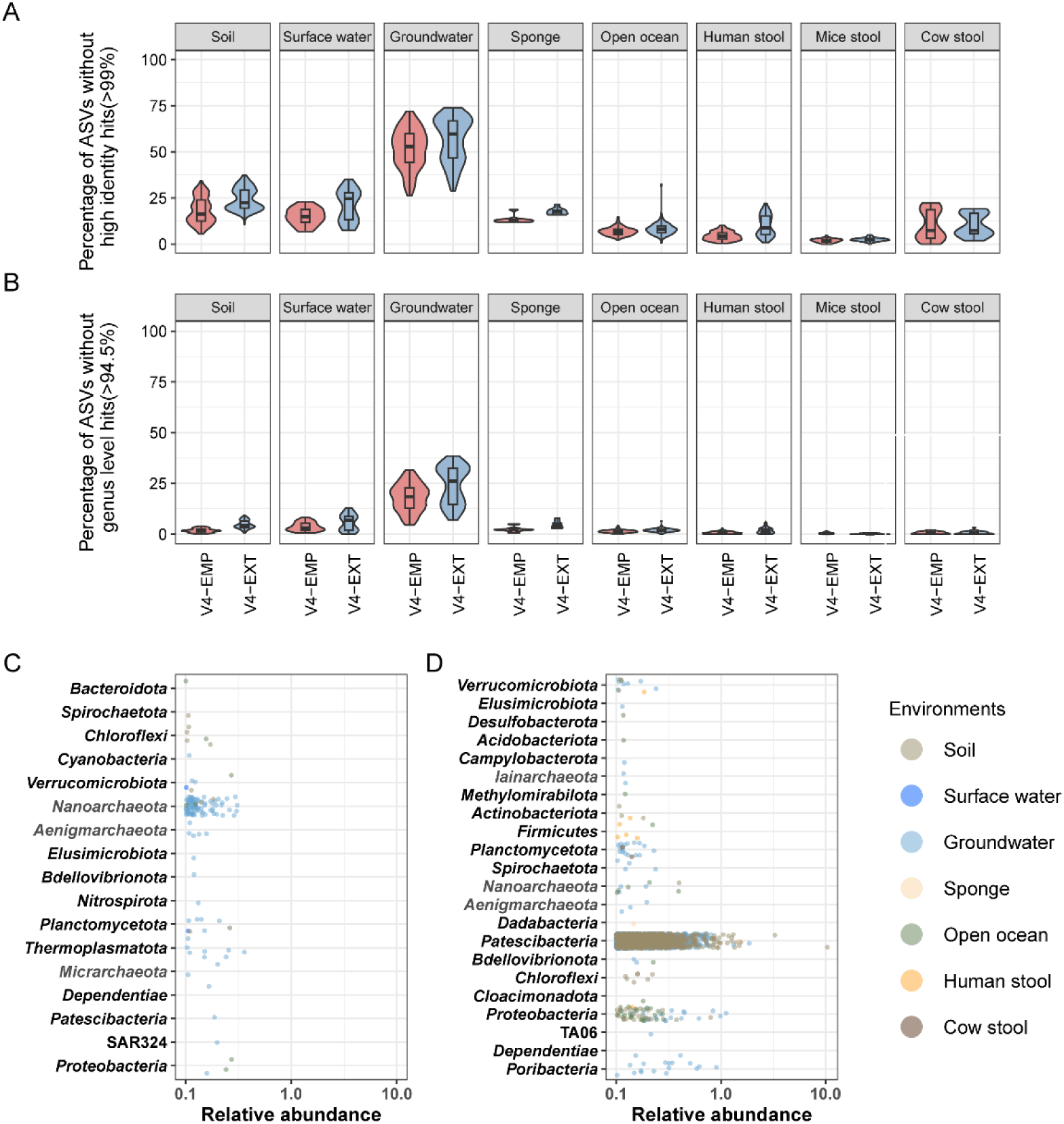
Comparison of ASV novelty generated by the V4-EMP (red) and V4-EXT (blue) primer pairs. The percentage of ASVs in each sample type A) without high-identity (>99%) matches and B) without genus-level identity (>94.5%) matches to the SILVA r138.2 database. The phylum-level classification of ASVs without genus-level matches in the SILVA v138.2 database in C) the V4-EMP and D) the V4-EXT dataset. Each dot represents an ASV in a sample, and only ASVs with >0.1% relative abundance are shown. Relative abundance is displayed on the x-axis, on a logarithmic scale. Different sample types are depicted by dots of different colors. The phylum SAR324 clade (Marine group B) is indicated by SAR324. Archaeal phyla are shown in grey font, and bacterial phyla are shown in black font.

To examine unique novelty detected by either primer pair, we further examined the ASVs lacking genus-level matches in the SILVA database that were only detected in the V4-EMP or the V4-EXT datasets. We found 56,278 ASVs with a relative abundance >0.01% without a genus-level match in the SILVA database unique to the V4-EXT dataset, while the V4-EMP dataset contained only 14,744 unique genus-level novel ASVs (Figure S3). *Nanoarchaeota* ASVs comprise 63.5%(n= 9369) of these novel ASVs in the V4-EMP dataset, followed by *Patescibacteria* (n=976, 6.6%), *Proteobacteria* (n=727, 4.9%), and *Verrucomicrobiota* (n=678, 4.6%) ASVs. In the V4-EXT datasets, 88.4% (n=49754) of the genus-level novel ASVs were classified as *Patescibacteria*, 2.2% (n=1260) as *Verrucomicrobiota*, 2.0% (n=1137) as *Proteobacteria*, 1.4% (n=783) as *Iainarchaeota*, and 0.7% (n=446) as *Nanoarchaeota* (Figure S3). The overwhelming majority of genus-level novel ASVs in both datasets (98.6% and 99.3% in V4-EXT and V4-EMP datasets, respectively) were extremely rare and displayed a cumulative relative abundance <0.1%.

Notably, among the ASVs without genus-level matches and with a relative abundance above 0.1%, only 2% ASVs (n=110) were unique to the V4-EMP datasets, most of which were classified as *Nanoarchaeota* (n=74, 67.3% of novel ASVs). They primarily originated from groundwater samples and showed individual relative abundances <0.3% (Figure 3C). The V4-EMP primer pair also uniquely detected 29 novel genus-level bacterial ASVs, which were distributed among 13 different phyla (Figure 3C). The V4-EXP primer pair detected 7 times more genus-level unassigned ASVs (n=792). The majority were identified as *Patescibacteria* (n=704, 88.9%), and originated from soil and groundwater ecosystems, highlighting substantial undiscovered diversity within this bacterial phylum, not captured by the V4-EMP primers. Additionally, we identified numerous novel, relatively highly abundant ASVs belonging to the order *Rickettsiales* (*Alphaproteobacteria*), detected across freshwater, soil, and ocean water column samples, as well as novel *Poribacteria* ASVs from freshwater environments (Figure 3D).

## Discussion

Here, we present an additional modification to a recently published primer pair [9] targeting the V4 region of bacterial and archaeal 16S rRNA genes (V4-EXT), with an even greater overall *in silico* coverage. We experimentally compared the performance of the V4-EXT primer pair to the V4-EMP primers, which are the most commonly used 16S rRNA gene-targeted primers across many fields of ecological research [9] using sequence datasets generated with both primer pairs from 938 samples from different ecosystems.

*In silico* analysis showed that the V4-EXT primer pair should particularly improve the ability to detect *Patescibacteria, Chloroflexi*, and *Iainarchaeota*. Consistently, in empirical datasets, the relative abundance of *Patescibacteria* was significantly higher across all sample types in V4-EXT datasets, except in human stool samples, in which these bacteria are nearly absent (Figure 2). Also, marked genus and species novelty amongst the *Patescibacteria* was detected when V4-EXT primers were applied (Figure 3). Significant and drastic differences in community composition data between datasets were detected particularly in high richness, high diversity samples from soil and groundwater environments. Several recent metagenomic surveys support our findings that groundwater and soils are major reservoirs of untapped prokaryotic diversity, especially regarding Patescibacteria and the archaeal DPANN superphylum [13,14]. In groundwater samples, where *Patescibacteria* accounted for on average >35% of all amplicon sequences in V4-EXT datasets, their relative abundance is severely underestimated with the V4-EMP primers (6.4%). The V4-EXT results align closer with observations from groundwater metagenomes, where *Patescibactria* are reported to be the dominant phylum with relative abundances ranging from 10–40% [14,15], and are increasingly recognized as abundant core community members. *Patescibacteria* are characterized by ultra-small cells and streamlined genomes with severely reduced biosynthetic and regulatory capacities, suggesting strong reliance on external sources for metabolites and interactions with partner organisms [16,17]. Studies indicate that specific *Patescibacteria* co-occur with specific other bacterial or archaeal taxa, forming symbiotic interactions, usually as parasitic epibionts [18,19].

Similarly, members of the archaeal superphylum DPANN, which, amongst others includes the phyla *Nanoarchaeota, Aenigmarchaeota, Woesearchaeota, Micrarchaeota*, and *Iainarchaeota*, are small cells with small genomes and limited metabolic capacities [20]. Like the *Patescibacteria*, many DPANN archaea are obligate episymbionts or parasites, physically attaching to host cells, such as *Thermoproteota* [21]. Metagenomic data suggest they can play a role in carbon, sulfur, and nitrogen cycling, though the extent of their involvement and influence on these elemental cycles remains poorly constrained [22]. The V4-EXT primers showed a significantly and markedly improved coverage of *Iainarchaeota* (Table S1), resulting in their significantly improved detection in groundwater samples (Figure 2). Small, enigmatic archaeal cells [16] have been shown to comprise up to 25% of groundwater microbial communities, with relative abundances of individual phyla being highly dynamic and patchy [14]. Thus, an inclusive detection of all DPANN phyla is important for accurate interpretation of their environmental significance and distribution. However, it is important to acknowledge that in our dataset, *Nanoarchaeota* still manyfold outnumber *Iainarchaeota* (Figure 2), and despite equal *in silico* coverage, the V4-EXT primer detected a significantly lower relative abundance and sequence novelty within the *Nanoarchaeota* than the V4-EMP primers (Figure 2, Figure S2). We were in this study not able to resolve if this discrepancy between *in silico* coverage and experimental detection of *Nanoarchaeota* is due to a real difference in primer performance or is based on community shifts due to the improved detection of other taxa, and could be ameliorated by increased per-sample sequencing depth.

We also found that the V4-EXT primers more effectively detected *Poribacteria* and *Chloroflexi* in sponge samples. These results align with global surveys of sponge microbiomes, conducted with a V3-V4 region-targeted 16S rRNA primer pair, which showed that both *Chloroflexi* and *Poribacteria* are enriched in marine sponges [23]. In metagenomic datasets from the Red Sea sponge *Hyrtios erectus, Poribacteria* accounted for ∼7% and Chloroflexi for 10.7% of the microbial community [24]. In a deep-sea sponge study, *Poribacteria* and Chloroflexi were identified to be among the most transcriptionally active phyla, contributing 17% and 12% of microbial transcripts, respectively, second only to *Proteobacteria* (23.7%) [25]. Beyond sponges, *Chloroflexi* are also known to be abundant in some soils and sediments, representing 9–16% of the bacterial community in, for example, river sediment samples from Rifle, Colorado [26], and reaching up to 60% relative abundance in deep marine sediments [27] and global soil diversity surveys [13].

Notably, several, mainly rare (<1% relative abundance) phyla were found to be statistically significantly less abundant in datasets generated with the V4-EXT (Figure S2), despite equal or better *in silico* coverage of the corresponding phyla (Table S1). These shifts are more likely observed due to the improved detection and related increased abundance of other taxa, rather than due to primer biases as illustrated by the minimal unique sequence novelty detected in the V4-EMP dataset (Figure 3).

While no difference in co-amplification of chloroplast 16S rRNA gene and eukaryotic 18S rRNA gene sequences was observed between primer pairs across sample types, mitochondrial 16S rRNA gene sequences were more strongly affected by primer choice, with the V4-EXT primers resulting in approximately three times more mitochondrial reads (Table S3). Still, the highest proportion, observed in soil samples, was only about 10% and is not high enough to prohibit the use of V4-EXT in any of the here tested source environment types, including directly host-derived samples (sponge and fecal samples).

In summary, we here present the V4-EXT primer pair - an improved successor to V4-CPR [9] and revision of the well-established and widely used V4-EMP primers [7,8], with equal or improved *in silico* coverage across all bacterial and archaeal lineages. We furthermore demonstrate using a large and diverse collection of samples that the application of V4-EXT has no detrimental drawbacks, but drastically alters microbial community compositions recovered from various environmental habitats, where previously poorly covered taxa like the *Patescibacteria* can account for up to on average 35% of all reads.

## Methods

### Sample collections and amplicon sequencing

We analyzed 938 samples from human and animal stool, groundwater, surface water, ocean, sponge, and soil collected in the framework of 12 independent research projects (Table S2). DNA extraction for soil samples from projects JMF-2401-02, JMF-2401-04, and JMF-2401-05 was performed using ∼250 mg soil (wet weight) per sample and the Qiagen DNeasy PowerSoil Pro Kit with the addition of 15% of a 20% skimmed milk solution to the CD1 buffer. Samples from project JMF-2210-08 and JMF-2212-20 were extracted using the MP Biomedicals FastDNA™ SPIN Kit for Soil from ∼400 mg soil (wet weight) following the respective manufacturer’s instructions. DNA from human (∼260 mg stool per sample; JMF-2206-10; JMF-2212-18), mouse (one fecal pellet per sample; JMF-2303-06, cow, and calf stool (∼250 mg stool per sample; JMF-2305-08) samples were extracted using the QIAamp Fast DNA Stool Mini Kit following the manufacturer’s instructions. DNA from open ocean surface samples was extracted using the Monarch Genomic DNA Purification kit (New England Biolabs, Ipswich, US) and the QIAwave Blood and Tissue kit (QIAGEN GmbH, Hilden, Germany). Surface water and groundwater samples were extracted using a phenol-chloroform-isoamyl alcohol extraction [28]. Water was filled into sterile canisters at the sampling site, and cells were collected on a Sterivex filter (0.2 µm pore size) until clogging (∼2 L for surface water, and ∼10 L for groundwater). DNA from snap-frozen shallow water marine sponge tissue samples (JMF-2407-25) was extracted using the DNase PowerSoil Pro kit (QIAGEN GmbH, Hilden Germany) following the manufacturer’s instruction. Sponge samples were collected under the permit G21/38062.1 and G38062.1 issued by the Great Barrier Reef Marine Park Authority. Amplification, barcoding, normalisation, pooling, and sequencing library preparation for all samples was performed using a two-step PCR protocol described in [29]. PCR mastermixes and primer concentration for both primers, as well as cycling conditions for the V4-EMP primers were previously optimized for this protocol and kept as published [29] after a 3 min initial denaturation at 94°C, 30 cycles of denaturation were performed for 45 sec at 94°C, annealing for 60 sec at 52°C, and elongation for 90 at 72°C, before a final elongation for 10 min at 72°C. For the V4-EXT primers (forward/reverse: GTGYCAGMMGBNKCGGTVA/RGACTAMNVRGGTHTCTAAT) after a 3 min initial denaturation at 95°C, 35 cycles of denaturation were performed for 40 sec at 95°C, primer annealing for 120 sec at 55°C, and elongation for 60 sec at 72°C, and a final elongation for 7 min at 72°C.For both primer pairs, first step PCRs were performed in 50 L reaction volume, with the DreamTaq Green PCR Master Mix (ThermoFisher), 0.25 μmol/L of forward and reverse primers each, and 5 μl DNA template. First step PCR products were then cleaned up and normalised using the SequelPrep Normalization Kit (Invitrogen), and 10 pL of first-step PCR product was added as template to a 50 pL barcoding PCR reaction, again containing the DreamTaq PCR Master Mix (ThermoFisher), with 0.8 μmol L^−1^ of two unique barcoding primers [29]. The barcoding PCR conditions were initial denaturation at 94°C for 4 min; 7 cycles of denaturation for 30 sec at 94°C, annealing for 30 sec at 52°C, and elongation for 60 sec at 72°C; followed by a final elongation step at 72°C for 7 min. Resulting amplicons were purified, normalised, pooled, library prepped, and amplicon libraries were sequenced on the lllumina MiSeq Platform, using v3 chemistry and 600 cycles (2x300 bp) as described previously [29].

### Amplicon sequence data analysis

Forward reads were trimmed at 220 nt, and reverse reads were trimmed at 150 nt with expected errors of 2 for barcode and primer sequences, respectively. Amplicon sequence variants (ASVs) were inferred from quality-trimmed reads with DADA2 package version 1.20.0 using the default recommended workflow (https://f1000research.com/articles/5-1492). Chimeras were detected by removeBimeraDenovo function in the DADA2 workflow and removed before further analysis. To assure robust sequence novelty analysis, the absence of further chimeric sequences was confirmed by the usearch package with the-uchime3 denovo workflow [30]. ASVs were classified by the assignTaxonomy function of DADA2 with the SILVA r138.1 database taxonomy, with a confidence threshold of 0.5. The fraction of off-target amplicons (i.e., ASVs classified as chloroplast, mitochondria, and eukaryote) in each sample was quantified before they were removed for all downstream analysis. Amplification and sequencing were performed on 1513 samples and data from a total of 938 samples, each with more than 5000 reads in the V4-EXT and V4-EMP dataset after exclusion of off-target amplicons, were included for analyses. Alpha diversity was summarized by the function amp_alphadiv using the ampvis2 package with rarefy=5000. Linear regression analysis was performed using the lm() function in R 4.1.2 [31]. To test for statistically significant differences between datasets, paired Wilcoxon tests were performed with the wilcox.test() function with R 4.1.2 [31]. Sequence novelty analysis was performed by blasting ASV sequences against the SILVA database r138.2 with evalue=1e-5. Hits with identity >= 99 or >= 94.5 and hit length > 200 bp were used for the ASV novelty analysis.

## Data availability

The 16S rRNA gene amplicon sequence data and associated metadata are available in the NCBI Sequences Read Archive (SRA) under the following BioProject IDs: PRJNA1366826 (JMF-2206-10), PRJNA1363873 (JMF-2210-08), PRJNA1366128 (JMF-2212-18), PRJNA1363863 (JMF-2212-20), PRJNA1226173 (JMF-2303-06), PRJNA1366369 (JMF-2305-08), PRJNA1366129 (JMF-2301-09), PRJNA1366130 (JMF-2302-05), PRJNA1367499 (JMF-2311-06), PRJNA1366290 (JMF-2401-02), PRJNA1366145 (JMF-2401-04), PRJNA1366146 (JMF-2401-05), PRJNA1367501/PRJNA1369818 (JMF-2407-25).

## Acknowledgements

This study was supported by the Wittgenstein Award of the Austrian Science Fund (FWF) [Z383-B] awarded to M.W., the Austrian Science Fund (FWF) Cluster of Excellence “Microbiomes drive Planetary Health’ (10.55776; COE 7; I.C.W., S.T., D.S., M.W., P.P.), the Austrian Science Fund (FWF) Principal Investigator Project 10.55776; P34775 (I.C.W.), the Austrian Science Fund (FWF) Hertha Finnberg Program Grant T 1218 (B.G.), the Austrian Science Fund (FWF) “Zukunftskolleg”/YIRG Project “playNICE” ZK-74B (P.P.), the Research Foundation Flanders (FWO) G0A4821N, G000821N and 1174925N (L.S.), and the Special Research Fund of Ghent University, through the Methusalem mandate of P.D (project no. BOF.MET.2021.0004.01). The processing of the amplicon sequencing data presented in work has been performed using the Life Science Compute Cluster (LiSC) of the University of Vienna. We additionally thank Gudrun Kohl, Sara Malinowski, Julia Ramesmayer, and Jasmin Schwarz for their help with DNA extraction and amplicon sequencing.

## Abbreviations

rRNA: Ribosomal ribonucleic acid
V4-EXT: V4 extend primer pair developed in this study
V4-EMP: V4 primer pair used in Earth Microbiome Project
V4-CPR: V4 primer pair developed for enhancing coverage of *Patescibacteria*
WWTPs: Wastewater treatment plants
ASVs: Amplicon sequence variants

## Notes

### Competing Interest Statement

The authors have declared no competing interest.

### Summary of Updates

A wrong number of PCR conditions was corrected.

