## Supplementary Figures 1,2,3 for "Revised 16S rRNA V4 hypervariable region targeting primers enhance detection of *Patescibacteria* and other lineages across diverse environments"

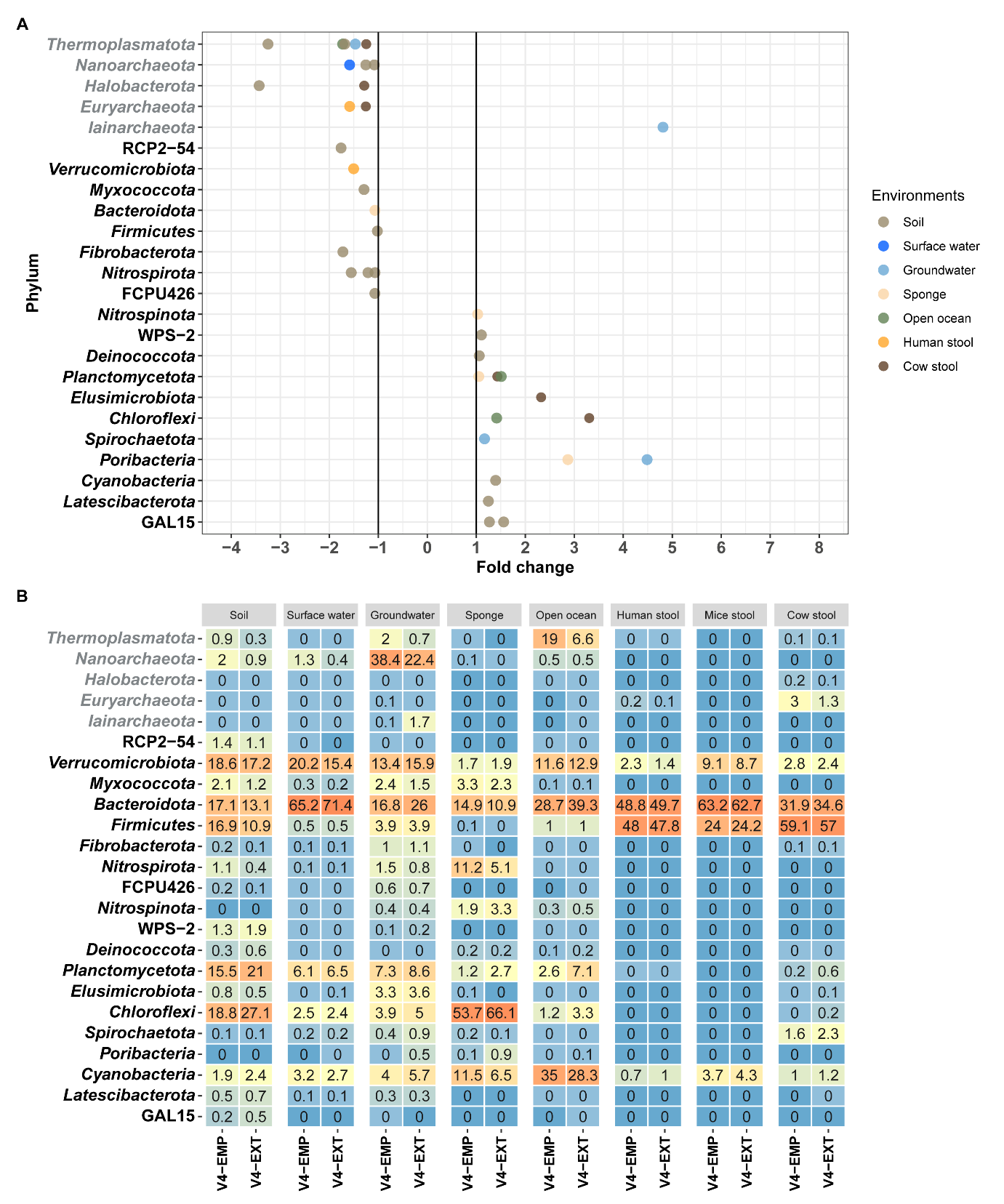

**Figure S1.** A) Fold change in the relative abundance of each phylum in datasets, grouped by source environment, generated using the V4-EXT versus the V4-EMP primes, after reads classified as *Patescibacteria* were removed. Phyla with a mean abundance >0.1% in any dataset are depicted. Phyla with a mean relative abundance of >0.1% for which mean relative abundance differences between datasets were above twofold are included. Fold change >1 indicates the phylum has a higher mean relative abundance in the V4-EXT dataset, and fold change <-1 indicates the phylum has a higher mean relative abundance in the V4-EMP dataset. B) Phyla level average relative abundance heatmap across different types of samples grouped by source environment, generated using the V4-EXT versus the V4-EMP primers, after reads classified as *Patescibacteria* were removed. Phyla with a mean relative abundance of >0.1% in either dataset are presented. Archaeal phyla are shown in grey font, and bacterial phyla are shown in black font.

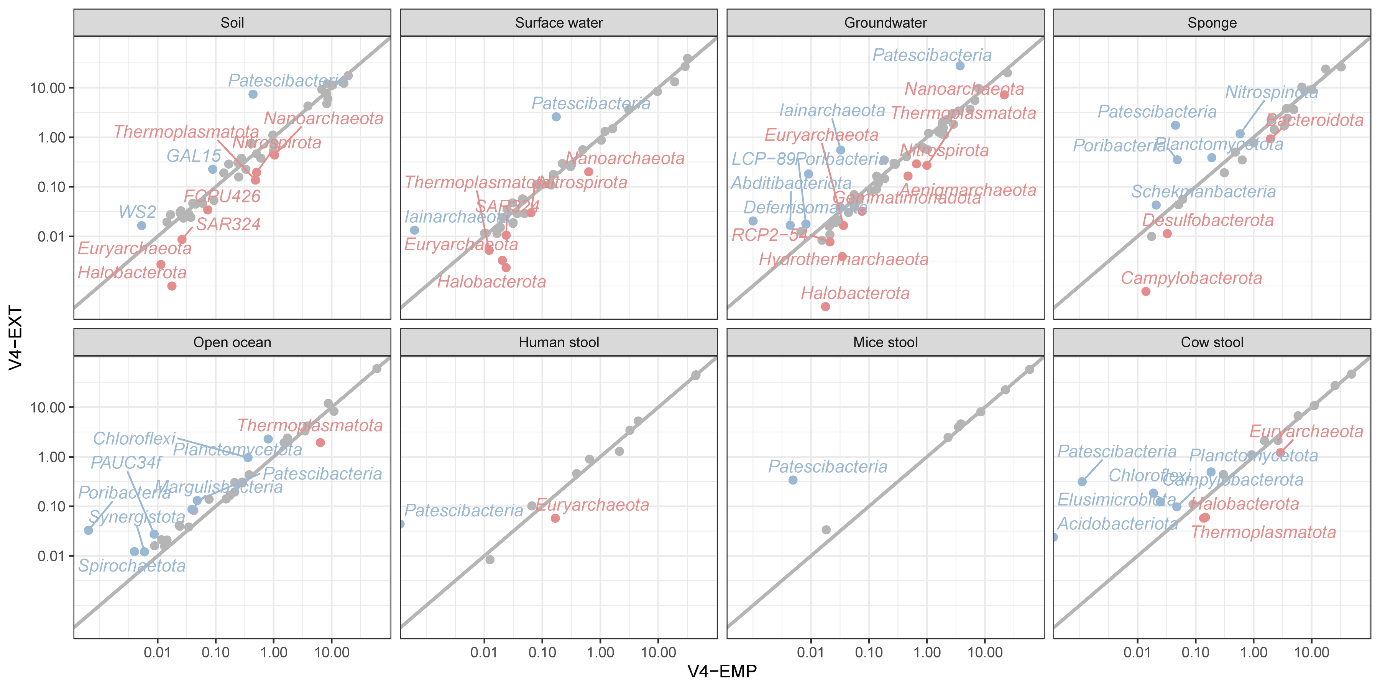

**Figure S2.** Comparison of the mean relative abundance (%) of phyla across different sample types in V4-EXT and V4-EMP datasets. Phyla with less than 0.01% average relative abundance in the V-EMP4 or V4-EXT datasets are not depicted. Grey: phyla with less than twofold difference in relative abundance between datasets. Blue: phyla overrepresented by at least twofold with V4-EXT. Red: phyla overrepresented by at least twofold with V4-EMP. The phylum name SAR324 clade (Marine group B) is indicated as SAR324.
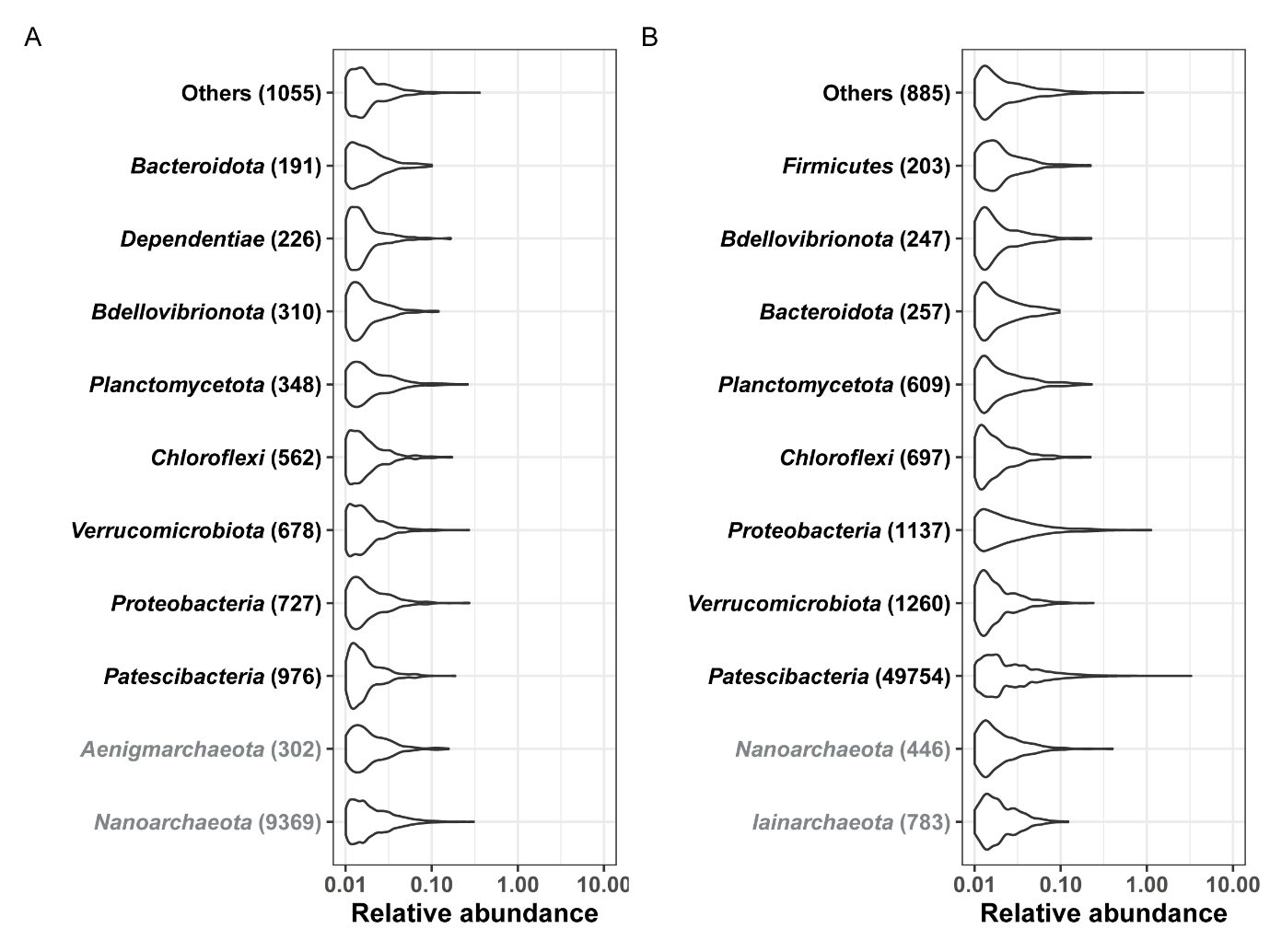

**Figure S3.** Phylum-level classification of ASVs (relative abundance>0.01%) without genus-level hits in the SILVA v138.2 database, generated by A) the V4-EMP B) V4-EXT primer pair. Each dot represents an observation of an ASV in a sample, and only observations with >0.01% relative abundance are shown. Per sample relative abundance is displayed on the x-axis, on a logarithmic scale. Top10 phyla are shown in the plot, while other phyla are collapsed in others. Archaeal phyla are shown in grey font, and bacteria phyla are shown in black font.

**Supplementary table legends**

**Table S1.** Phyla-specific in silico coverage of the three analyzed 16S rRNA gene V4 region primer pair versions using the SILVA database (SILVA_138.2_SSURef_NR99) as reference. Numbers indicate the percentage of sequences from each phylum in the SILVA database covered without mismatch or with one mismatch by the V4-EMP, V4-CPR, and V4-EXT primers. Archaeal phyla are shown in grey font, and bacterial phyla are shown in black font.

**Table S2.** Summary of Sample JMF IDs and Dataset JMFS IDs, as well as their source habitats, used in this study.

**Table S3.** The percentage of sequences classified as chloroplast, mitochondrial, or eukaryotic 16S or 18S rRNA genes in the V4-EMP and V4-EXT datasets obtained from different sample types. The grey background and bold font indicates that a significant difference (pairwise Wilcoxon test, p<0.05) was detected between the V4-EMP and V4-EXT datasets.

**Table S4**. Fold changes in the relative abundances of each phylum in different datasets generated by the V4-EMP and V4-EXT primer pair originally and after Patescibacterial sequenced were removed.
